# Cholinergic signalling at the body wall neuromuscular junction couples to distal inhibition of feeding in *C. elegans*

**DOI:** 10.1101/2021.02.12.430967

**Authors:** Patricia G. Izquierdo, Thibana Thisainathan, James H. Atkins, Christian J. Lewis, John E.H. Tattersall, A. Christopher Green, Lindy Holden-Dye, Vincent O’Connor

**Affiliations:** Biological Sciences, Institute for Life Sciences, University of Southampton, Southampton, United Kingdom; Defence Science and Technology Laboratory Porton Down, Salisbury, United Kingdom

**Keywords:** Acetylcholinesterase inhibitor, levamisole, pharyngeal pumping, inter-tissue communication, neuromuscular junction, tissue specific rescue

## Abstract

Complex biological functions within organisms are frequently orchestrated by systemic communication between tissues. In the model organism *C. elegans*, the pharyngeal and body wall neuromuscular junctions are two discrete structures that control feeding and locomotion, respectively. These distinct tissues are controlled by separate, well-defined neural circuits. Nonetheless, the emergent behaviours, feeding and locomotion, are coordinated to guarantee the efficiency of food intake. We show that pharmacological hyperactivation of cholinergic transmission at the body wall muscle reduces the rate of pumping behaviour. This was evidenced by a systematic screening of the cholinesterase inhibitor aldicarb’s effect on the rate of pharyngeal pumping on food in mutant worms. The screening revealed that the key determinant of the inhibitory effect of aldicarb on pharyngeal pumping is the L-type nicotinic acetylcholine receptor expressed in body wall muscle. This idea was reinforced by the observation that selective hyperstimulation of the body wall muscle L-type receptor by the agonist levamisole inhibited pumping. Overall, our results reveal that body wall cholinergic transmission controls locomotion and simultaneously couples a distal inhibition of feeding.

## Introduction

The communication between tissues has an important role in physiological processes in health, disease and stress conditions (1,2). The communication between the skeletal muscle, liver and adipose tissue during exercise is a clear example of tissue communication to control energy metabolism and insulin sensitivity in that particular stress condition (3). The imbalance of that communication causes an increase of energy expenditure associated with chronic disease or cachexia (4). This complex process is conserved from invertebrates to mammals and can be modelled in simpler organisms such as the fruit fly *Drosophila melanogaster* (5–7).

The survival of the model organism *Caenorhabditis elegans* depends on two essential behaviours, feeding and locomotion, as well as the ability to modify them according to the environmental cues. The external presence of food is a potent stimulus that modulates the rate of feeding by increasing pharyngeal movements and the intake of the external bacterial food source (8–10). Interestingly, nematodes can modify their locomotory patterns according to food availability, changing between dwelling and roaming (11,12). In this sense, the quality and quantity of food ingested emerges as an important environmental factor to modulate the worm’s motility that is directly controlled by the neuromuscular body wall (12,13). Similarly, mechanical stimulation or the optogenetic silencing of the body wall musculature reduces the feeding rate of *C. elegans* that is otherwise directly governed by the neuromuscular transmission of cholinergic transmission at the pharynx (14–16). This supports the notion that locomotory function might provide additional pathways that contribute to regulation of the pharyngeal circuits that control feeding.

Morphologically, the pharynx is divided into three different parts: the corpus, the isthmus and the terminal bulb. Two pharyngeal movements are responsible for the transport of bacteria by ingestion via the buccal cavity to the intestines (17–19). The coordinated contraction-relaxation cycle of the corpus and the terminal bulb causes the opening of the lumen in these parts of the pharynx that results in the aspiration of bacteria into the cavity. This rhythmic movement is named pharyngeal pumping and is caused by the release of acetylcholine and glutamate from the MC and M3 pharyngeal neurons that contract and relax the pharyngeal muscle, respectively (20,21). The bacteria accumulated in the corpus during the pumping are directed to the terminal bulb by the progressive wavelike contractions of the muscles in the isthmus. This peristalsis movement depends on acetylcholine release from the pharyngeal motor neuron M4 (22,23), one of the 20 neurons that more widely supports regulation of feeding. This circuit is isolated from the rest of the animal by a basal lamina (17). A single synaptic connection is described between the pharynx and the rest of the animal (I2-RIP). However, disruption of this connection does not cause any defect in the feeding phenotype, indicating that this anatomical route does not mediate a strong determinant of the pharyngeal modulation of feeding (19). This points to a more important role of neuromodulatory signalling via biogenic amines and peptides, involving volume transmission (10).

In comparison to the body wall muscle, less is known about the molecular composition of the transmitter functions that control the pharyngeal neuromuscular junction (NMJ) and the feeding phenotype. The glutamate-gated chloride channel AVR-15 acts postsynaptically in the pharyngeal muscle to sense glutamate released from the motor neuron M3 that cause muscle relaxation (20,24). EAT-2 is a cys-loop acetylcholine receptor subunit localized at the synapse between the pharyngeal MC motor neuron and the muscle at the corpus. It requires the auxiliary protein EAT-18 to allow EAT-2 essential function in initiating contraction (21,25). Mutations in *avr-15* and *eat-2* phenocopy the feeding behaviour of nematodes with M3 and MC ablated neurons respectively, highlighting the critical role these receptor components play (20,21,24). The feeding phenotype additionally requires the release of acetylcholine by the motor neuron M4, triggering isthmus peristalsis (23). However, there is not any mutation in an acetylcholine receptor that phenocopies the M4 ablation and the molecular determinants of this feeding critical function are unknown.

In order to better understand the pharyngeal NMJ of *C. elegans*, we performed a targeted screen of the pumping behaviour on food with defined molecular determinants of cholinergic transmission in the presence or absence of aldicarb. This acetylcholinesterase inhibitor has been previously used to induce paralysis of movement and this led to the discovery of molecular components at the body wall neuromuscular junctions in *C. elegans* (26,27). We found that the genes conferring significant drug sensitivity of the pharyngeal paralysis are surprisingly located in the body wall NMJ. Additionally, we describe how the stimulation of the L-type receptor at the body wall muscle by the specific agonist levamisole reduced the pharyngeal pumping rate. This indicates an unexpected communication between the body wall neuromuscular junction that controls locomotion and the distinct and physically separated pharyngeal circuit that controls feeding.

## Results

### The determinants that control pharyngeal function are distinct from the determinants that control pharyngeal sensitivity to aldicarb

In order to investigate the cholinergic regulation of the feeding behaviour, the well-characterized protocol of aldicarb-induced paralysis was performed (26,27). This assay has been extensively used to find molecular determinants at the NMJ of the body wall on the basis of resistance or hypersensitivity to paralysis of locomotion when nematodes are incubated on aldicarb-containing agar plates (26). We previously demonstrated that aldicarb and other cholinesterase inhibitors also cause a dose-dependent inhibition of the pharyngeal function (28). This inhibition was associated with the pharynx exhibiting hypercontraction observed by the opening of the lumen (28). This implied that aldicarb intoxication in the context of the whole organism would result from drug-induced hyperactivation of the cholinergic transmission. In this sense, we hypothesized that molecular determinants in the pharynx would confer resistance or hypersensitivity to aldicarb-induced paralysis of feeding behaviour.

We screened the pharyngeal function of different mutant worms in the presence or absence of the cholinesterase inhibitor aldicarb under the conditions we previously optimized (28). The results of this screening are listed in Table 1. It consisted of 53 mutant strains deficient in cholinergic and other neurotransmitters signalling. The components of the cholinergic pathway tested included alpha and non-alpha subunits of the acetylcholine-gated cation channel (29), subunits of the nematode selective acetylcholine-gated chloride channel (30), muscarinic acetylcholine receptors (31–34), acetylcholinesterases (35–37) and auxiliary proteins involved in the proper function of the cholinergic receptors (21,25,38–47) (Table 1). This analysis can be conveniently summarized by ascribing responses into three different groups of mutants regarding their pharyngeal pumping on food in the absence and presence of aldicarb (Fig. 1).

**Table 1:**
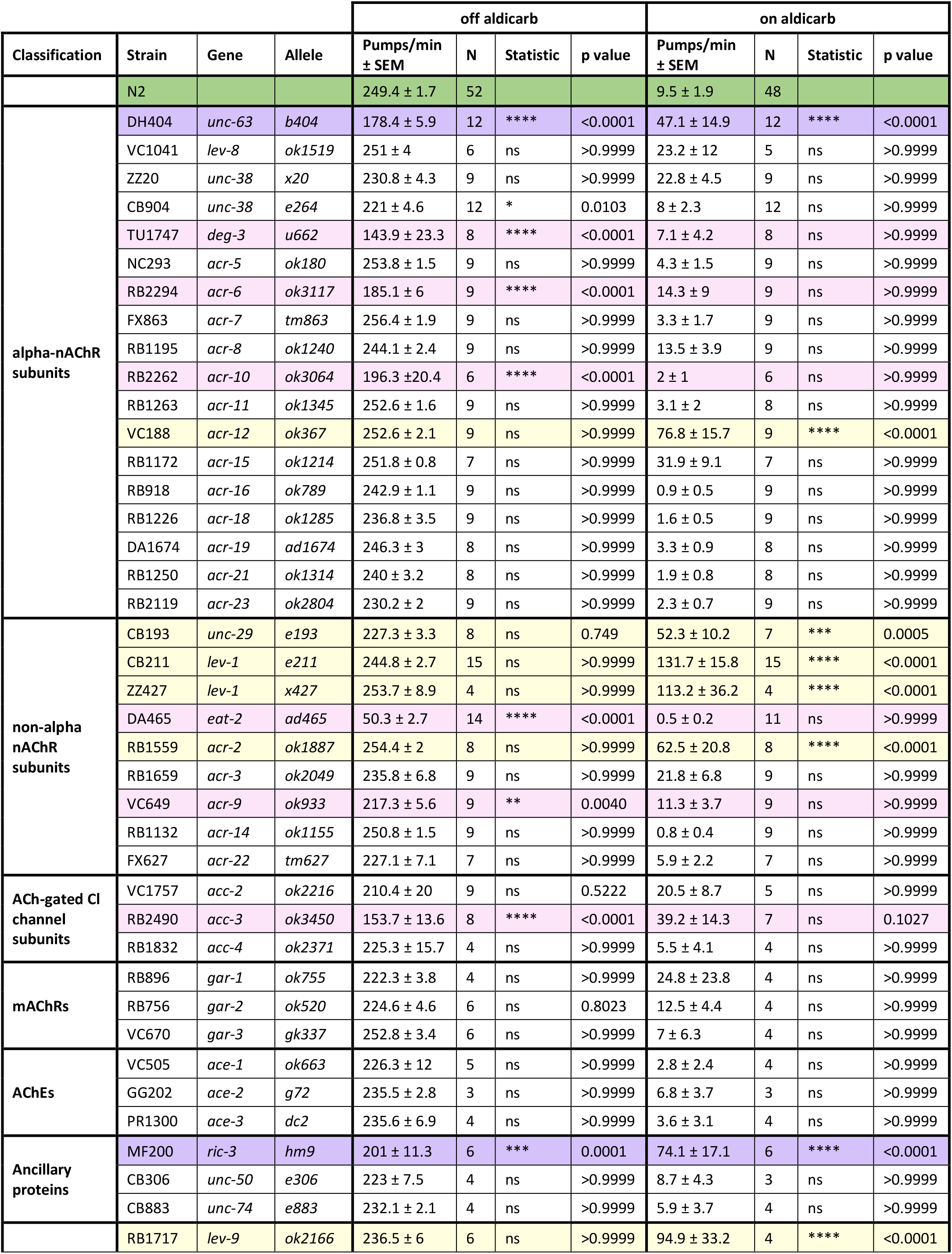
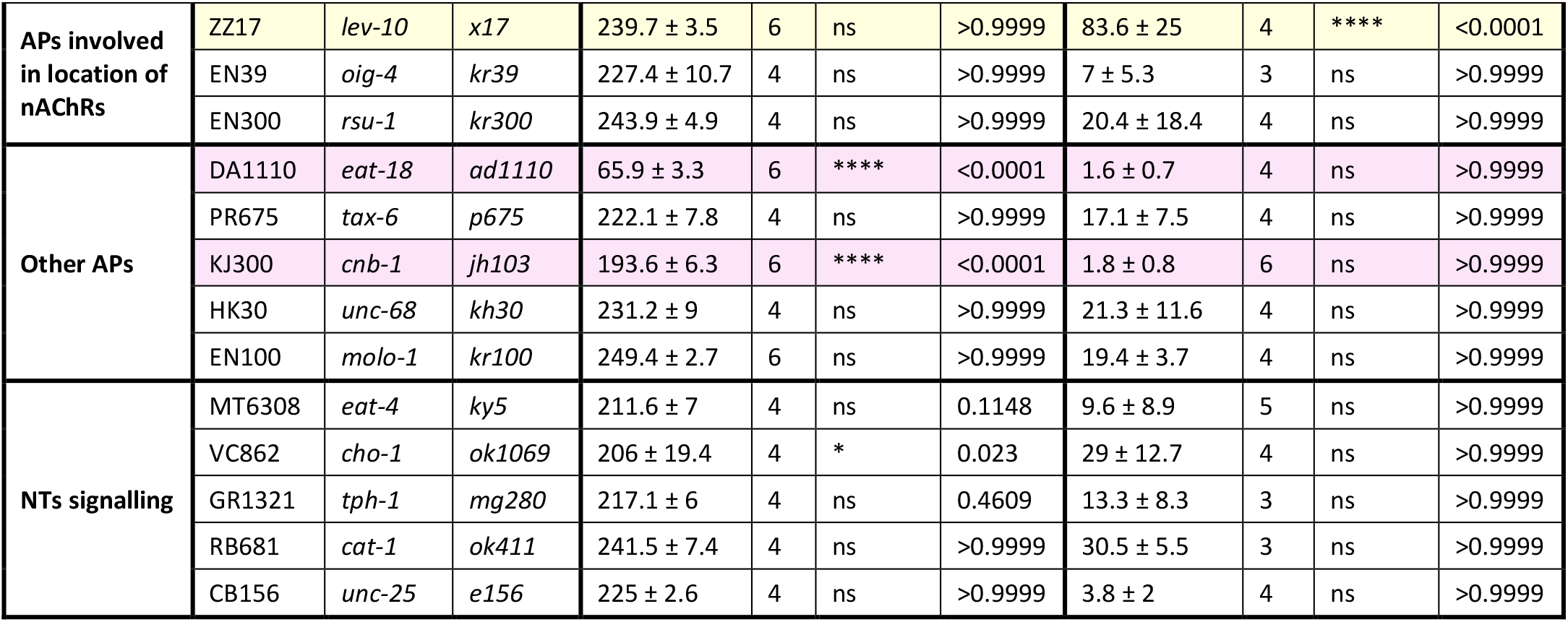
Pharyngeal pumping rate on food in the absence or presence of aldicarb. Synchronized L4+1 nematodes were incubated for 24 hours on seeded plates containing aldicarb or vehicle control before pumping rate per minute was quantified. Refer to Figure 1 for colour code. nAChR: acetylcholine-gated cation channel; mAChRs: muscarinic acetylcholine receptors; AChEs: acetylcholinesterases; APs: auxiliary proteins; NTs: neurotransmitters. Statistical analysis corresponds to the comparison between each strain and the N2 wild type control in each condition (off and on aldicarb). Data are shown as mean ± SEM. ^ns^p≥0.05; *p˂0.05; **p˂0.01; ***p<0.001; ****p<0.0001 by two-way ANOVA test followed by Bonferroni corrections.

**Figure 1:**
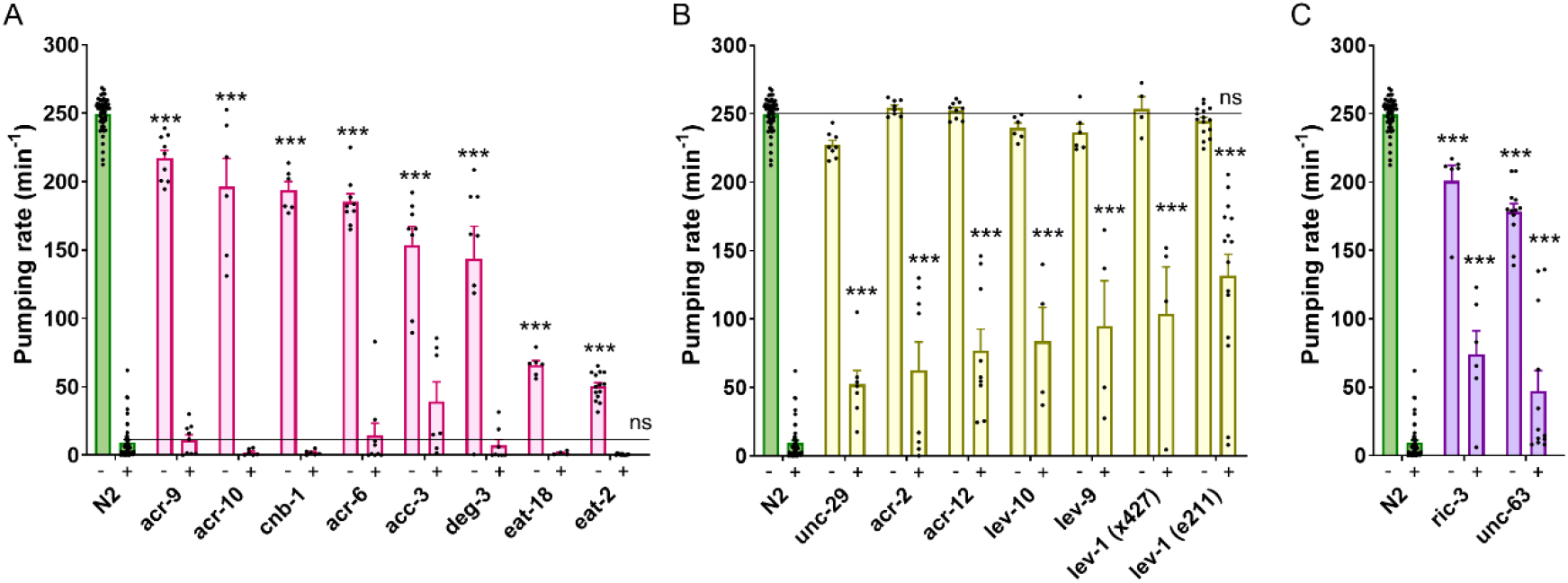
Molecular determinants that control pharyngeal function are distinct from the determinants that confer pharyngeal resistance to aldicarb. Pharyngeal pumping on food in the absence (−) or presence (+) of 500 μM aldicarb. A) Mutant nematodes deficient in the non-alpha acetylcholine-gated cation channel subunits ACR-9 and EAT-2; the alpha acetylcholine-gated cation channel subunits ACR-10, ACR-6 and DEG-3; the acetylcholine-gated chloride channel subunit ACC-3 and the acetylcholine receptor auxiliary proteins EAT-18 and CNB-1 exhibited a significant reduction of the pumping rate but a normal sensitivity to aldicarb compared to the wild type control. B) Mutant nematodes deficient in the non-alpha acetylcholine-gated cation channel subunits UNC-29, LEV-1 and ACR-2; the alpha acetylcholine-gated cation channel subunit ACR-12 and the acetylcholine receptor auxiliary proteins LEV-9 and LEV-10 exhibited normal pumping rate on food but significant resistant to aldicarb compared to the wild type worms. C) Nematodes deficient in the ancillary protein RIC-3 and the alpha acetylcholine-gated cation channel subunit UNC-63 presented both phenotypes, reduced pumping rate on food and resistance to aldicarb compared to the wild type control. Data are shown as mean + SEM of the pumping per minute. Statistical analysis corresponds to the same condition (absence or presence of aldicarb) between the N2 wild type and each mutant strains. ^ns^p>0.05; ***p<0.001 by two-way ANOVA test. Refer to Table 1 for N numbers and p values.

One class of mutants showed reduced pumping rate in the absence of aldicarb but wild type inhibition of pharyngeal pumping in the presence of the drug (Fig. 1A). This class of mutants clearly harbours an important contribution to feeding at physiological levels of cholinergic stimulation when nematodes are on food. However, despite this essential contribution, no resistance to the inhibition was observed when the cholinergic stimulation increased as a consequence of aldicarb exposure. According to this, we identified this class of mutants as “physiological determinants” of the feeding phenotype. This is exemplified by the *eat-2* and *eat-18* mutants. In addition, mutant nematodes deficient in the subunits of the acetylcholine-gated cation channel DEG-3, ACR-6, ACR-9 and ACR-10, the subunit of the acetylcholine-gated chloride channel ACC-3 and the calcineurin CNB-1 exhibited a similar pattern of reduced pumping when they are off drug and similar pharyngeal inhibition on aldicarb compared to the response shown by the N2 wild type (Fig. 1A).

The second class of mutants essentially described the opposite of the above. These mutants showed normal pumping rate on food relative to N2 but clear resistance to the aldicarb-induced inhibition of the pharyngeal function observed in the wild type treated worms (Fig. 1B). The effect of this second class of determinants in the pharyngeal function was only apparent when the cholinergic transmission was overstimulated beyond the physiological levels by preventing acetylcholine degradation due to acetylcholinesterase inhibition by aldicarb. Therefore, we named this group “pharmacological determinants” of feeding. These mutants included the well characterized subunits of the acetylcholine-gated cation channel UNC-29, ACR-2, ACR-12 and LEV-1, along with the auxiliary proteins LEV-9 and LEV-10 (Fig. 1B). Interestingly, all the pharmacological determinants of the pharyngeal function are essential determinants of the body wall NMJ that controls locomotion (41,42,48,49).

Finally, the third class of mutants showed clear deficiency in the two distinct contexts, namely pumping on food in the absence of drug, and resistance to aldicarb-induced inhibition of the pharyngeal function (Fig. 1C). Only two genes of those tested were encompassed in this group (Fig. 1C), the alpha subunit of the L-type receptor UNC-63 and the chaperone of the nicotinic receptors RIC-3 (50,51)

Overall, the results suggest an unexpected divergence in cholinergic determinants of the pharyngeal function. Some of these control the physiological transmission that underpins fast pumping rate on food and others the hypothesized aldicarb-dependent process that executes an inhibition of feeding in the presence of the drug.

### The pharyngeal function of *C. elegans* exposed to levamisole exhibits a complex dose- and time-dependent inhibition

The pharmacological determinants of the pharyngeal function highlighted in the previous screening with aldicarb (Fig. 1B) are known to underpin the mode of action of the nematode selective pharmacological agent levamisole. This drug acts as an agonist of the body wall muscle nicotinic L-type receptor causing a spastic paralysis, essential for its use as a nematicide (52–54). Although distinct in its mode of action, levamisole, like aldicarb intoxication, leads to a hyperstimulation of the cholinergic synapses in the nematodes (54).

Accordingly, we investigated the levamisole response in the pharyngeal circuit by quantifying the pharyngeal pumping of wild type worms to increasing concentrations of drug over time (Fig. 2). Interestingly, nematodes displayed a profound initial inhibition of pharyngeal pumping rate when they were placed on levamisole-containing plates (Fig 2A). This reduction of the pharyngeal pumping was observed after control worms recovered from the mechanical stimulation caused by the picking process, which inhibits the feeding temporally (Fig. 2A) (55). The IC_50_ value calculated after 10 minutes of incubation onto levamisole plates was 140.3 μM (Fig. 2B).

**Figure 2:**
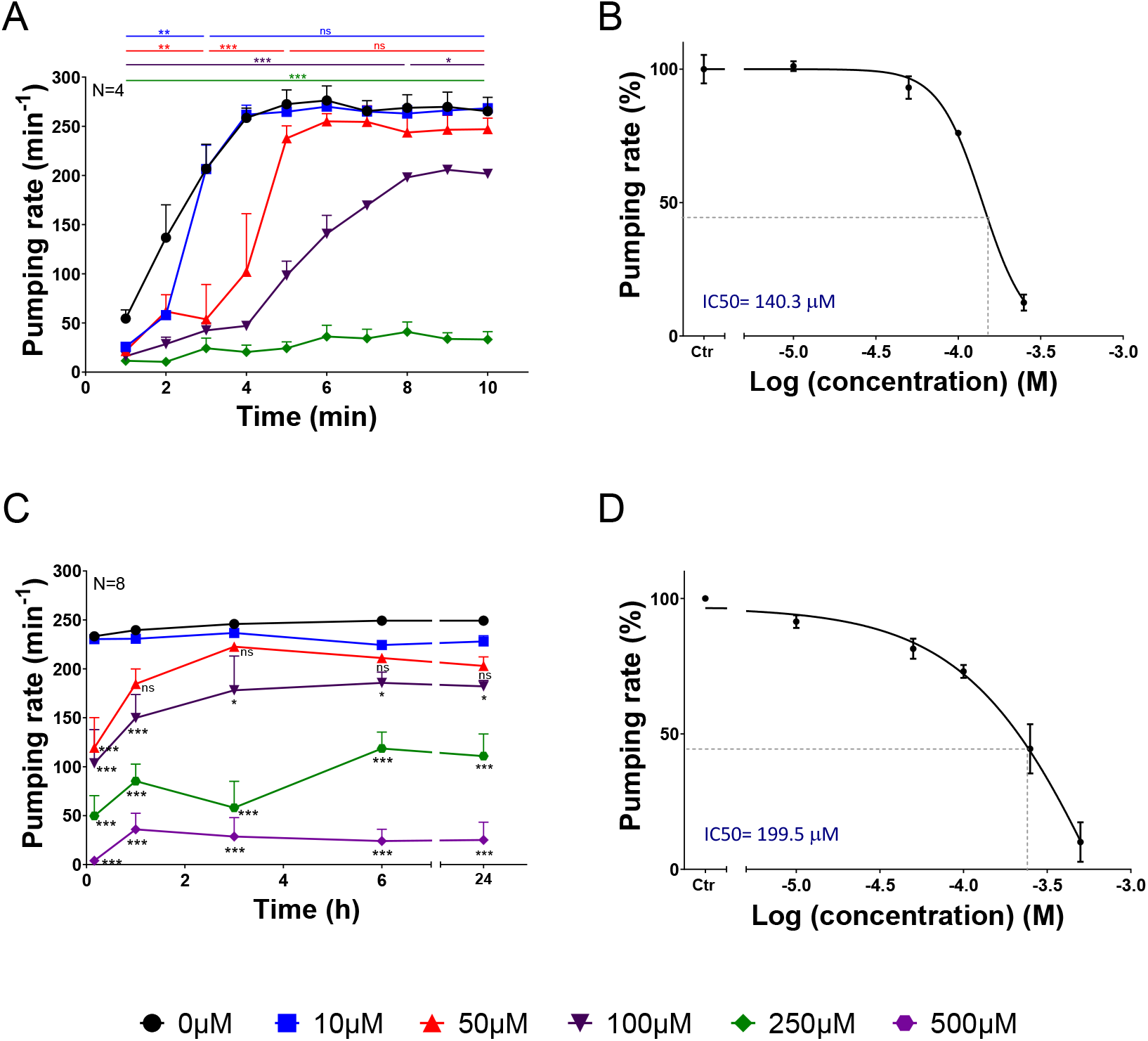
Pharyngeal function of *C. elegans* exposed to levamisole exhibited a complex concentration and time-dependent inhibition. A) Pharyngeal pumping was quantified for synchronized L4 + 1 nematodes immediately after transferring to either naïve or levamisole-containing plates. The initial picking-mediated inhibition of pumping (15) was recovered within 4 min. Nematodes picked onto levamisole-containing plates displayed a delayed dose-dependent recovery of the pharyngeal function after picking. Data are shown as mean + SEM of the pumping rate of 4 worms in 4 different experiments. B) IC50 value for pharyngeal inhibition by levamisole after 10 min of incubation. C) Pharyngeal pumping rate was quantified for synchronized L4 + 1 nematodes at different range of concentrations of levamisole over the time. An increased dose-dependent response was observed. Data are shown as mean + SEM of the pumping rate of 8 worms in 4 different experiments per dose. D) IC50 value for pharyngeal inhibition by levamisole after 24 hours of exposure. Data are shown as mean + SEM of the pumping per minute of 8 nematodes in 4 independent experiments. Statistical analysis corresponds to the comparison between each concentration and the non-drug control in each end-point time of incubation. ^ns^p>0.05; *p˂0.05; **p˂0.01; ***p<0.001 by two-way ANOVA test.

After this initial inhibition of the pharyngeal function by levamisole, wild type nematodes exhibited a partial recovery of the pumping rate over time at the lowest concentrations tested (10 μM and 50 μM) but the pumping was profoundly inhibited at the highest doses (250 μM and 500 μM) (Fig. 2C). This recovery of the pharyngeal function impacted on the IC_50_ value calculated, being 199.5 μM after 24 hours of incubation on levamisole-containing plates (Fig. 2D).

The fact that both levamisole and aldicarb inhibit pumping rate on wild type worms, and that inhibition is additionally reduced in *lev-1* deficient mutants for both drugs, support the hypothesis that the pharmacological hyperstimulation of the cholinergic system by either aldicarb or levamisole inhibits pharyngeal pumping by a common mechanism.

### The extra-pharyngeal nicotinic receptor subunit LEV-1 is a key determinant of levamisole inhibition of pharyngeal pumping

To more clearly resolve the molecular pathway through which the pharyngeal inhibition is mediated, we focused on the quantification of the pharyngeal function of *lev-1* deficient strains in the presence of levamisole (Fig. 3). The LEV-1 subunit was highlighted in our screen due to the selective contribution to aldicarb-induced modulation of the feeding (Fig. 1B). In addition, the LEV-1 subunit was originally identified as a determinant of body wall muscle sensitivity to levamisole (48).

**Figure 3:**
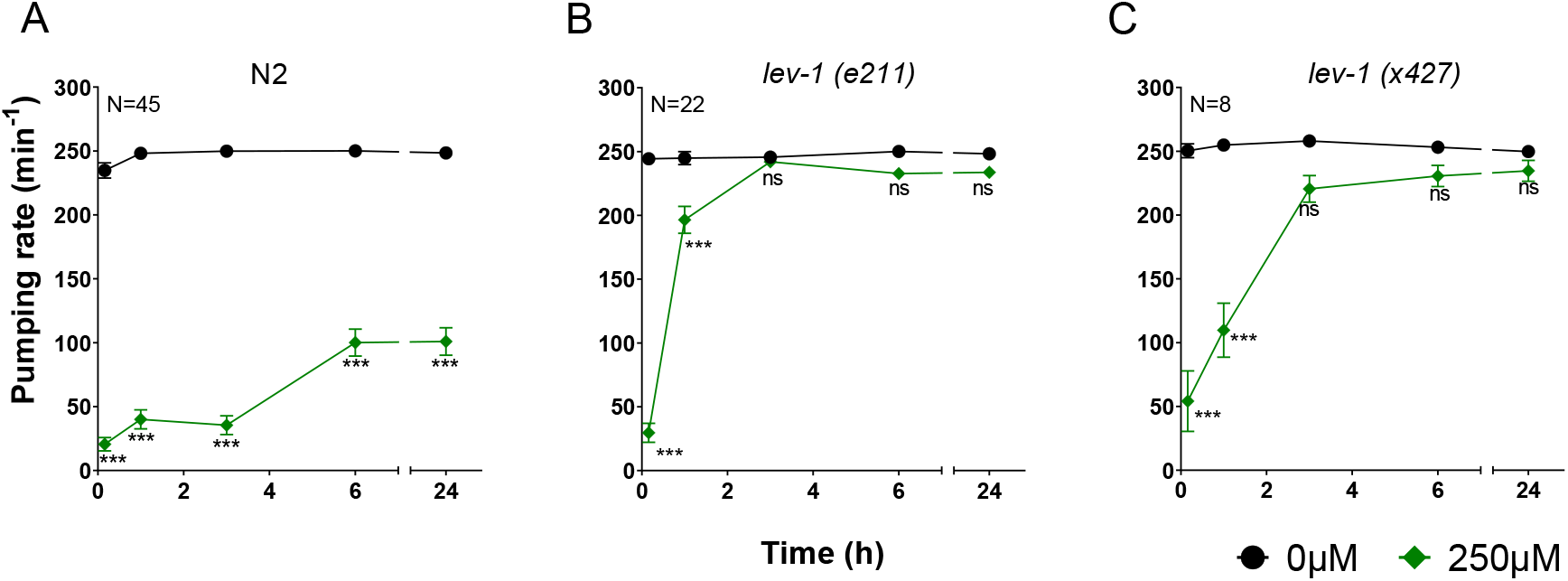
The non-alpha subunit LEV-1 of the heteromeric cholinergic receptor is responsible of the pharyngeal inhibition in the presence of levamisole at later end-point times. A) Pharyngeal pumping was measured in the presence or absence of 250 μM of levamisole. Data are shown as mean ± SEM of the pumping rate of 45 worms in at least 25 independent experiments. B) Pumping rate of CB211 *lev-1* strain in the presence or absence of 250 μM levamisole. Data are shown as mean ± SEM of 22 worms in at least 10 independent experiments. C) Pharyngeal pumping rate of ZZ427 *lev-1* mutant strain nematodes onto naïve or levamisole-containing plates. Data are shown as mean ± SEM of the pumping per minute of 8 nematodes in 4 independent experiments. Statistical analysis corresponds to the comparison between pumping rate on and off levamisole plates in each end-point time of incubation ^ns^p>0.05; ***p<0.001 by two-way ANOVA test.

Strains deficient in *lev-1* displayed a similar inhibition of the pharyngeal function as wild type worms after 10 minutes of incubation on 250 μM levamisole plates (Fig. 3). However, the pumping rate completely recovered over the time, being similar to the non-drug exposed nematodes after 3 hours of incubation (Fig. 3B and 3C). This phenotype was consistent in the two *lev-1* deficient strains tested, indicating the LEV-1 subunit of the nicotinic receptor is not responsible for the pharyngeal inhibition by levamisole at early exposure times, but its function is indeed required at the later exposure times.

These results highlight two distinct components of a complex response to worm intoxication by levamisole. While the rapid effect is independent of *lev-1*, the late sustained inhibition is clearly *lev-1* dependent. This points to an overlapping mechanism for the aldicarb and levamisole-induced inhibition of the pharynx at protracted intoxication conditions. This mechanism is mediated by a LEV-1 containing receptor.

Due to the pivotal role played by LEV-1, we sought to detail its expression beyond the well characterized body wall muscle expression (48). We first investigated the expression of *lev-1* in the pharyngeal circuit of *C. elegans* using existing GFP translational reporters previously used to address functional expression of *lev-1* (56). Our analysis of these strains supported previous descriptions about the location in the body wall muscle cells as well as nerve ring, dorsal and ventral nerve cord (56,57). However, previous investigations highlighted that some pharyngeal gene expression can be masked by overlying nerve ring expression (58). In order to address this, pharynxes from transgenic worms carrying transcriptional (AQ585) and translational (AQ749) GFP reporters (56) were isolated and imaged to test for fluorescent signal (Fig. 4A). We did not observe GFP fluorescence in any of the preparations, indicating LEV-1 does not occur in the isolated muscle or in its associated basal lamina embedded pharyngeal circuit (Fig. 4A). This was reinforced utilizing RT-PCR with primers designed to specifically amplify *lev-1* from mRNA extracted from the isolated pharynxes of wild type worms (Fig. 4B). The amplification of *myo-2* and *eat-2* were used as positive controls to demonstrate that the quality of mRNA was sufficient to amplify high and low copy number of transcripts. Selective amplification of *myo-3* was used as a negative control to show loss of body wall muscle transcripts as a consequence of anatomically isolating the pharynx. The mRNA of *lev-1* was not detected in the pharyngeal circuit of *C. elegans* worms (Fig. 4B).

**Figure 4:**
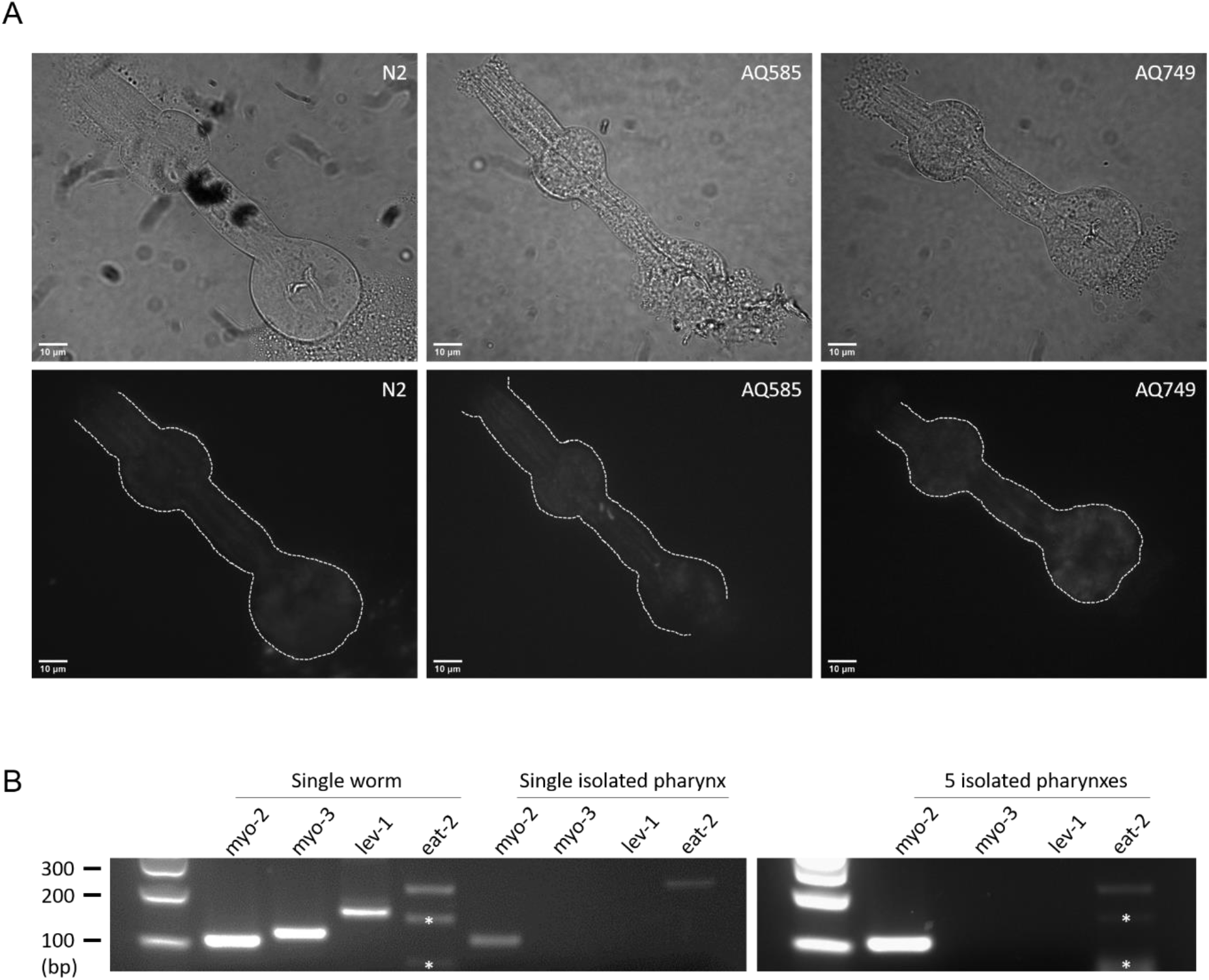
The determinants that control aldicarb and levamisole sensitivity in the pharynx are not expressed in the pharyngeal muscle of embedded circuits. A) Isolated pharynxes of transgenic lines expressing GFP under *lev-1* promoter in a transcriptional reporter (AQ585) or a translational reporter (AQ749) indicate there is not expression of LEV-1 in the pharynx. Dissected pharynxes of N2 wild type worms were used as negative control. Scale bar represents 10 μm. B) RT-PCR of N2 wild type isolated pharynxes demonstrates there is not expression of *lev-1* in the pharyngeal circuits. cDNA was reverse transcribed from a single worm, a single isolated pharynx or five isolated pharynxes RNA of wild type nematodes. PCR was performed using specific primers for *myo-2* and *eat-2* as positive control and *myo-3* as negative control of the expression in the pharynx. White asterisks correspond to unspecific amplification.

Taken together, these results indicate that the major pharmacological determinant of the drug-induced pharyngeal inhibition phenotype exerts its function outside the pharyngeal circuit.

### LEV-1 is required in the body wall muscle to mediate levamisole inhibition of pharyngeal pumping

In view of the significance of LEV-1 in the body wall NMJ, we investigated tissue specific rescue of LEV-1 at the musculature controlling the locomotion. For this, we generated transgenic lines of *lev-1* deficient nematodes expressing the wild type cDNA version of the gene under the control of either *lev-1* or *myo-3* promoter. This experiment was replicated with two distinct *lev-1* deficient mutant strains (Fig. 5).

**Figure 5:**
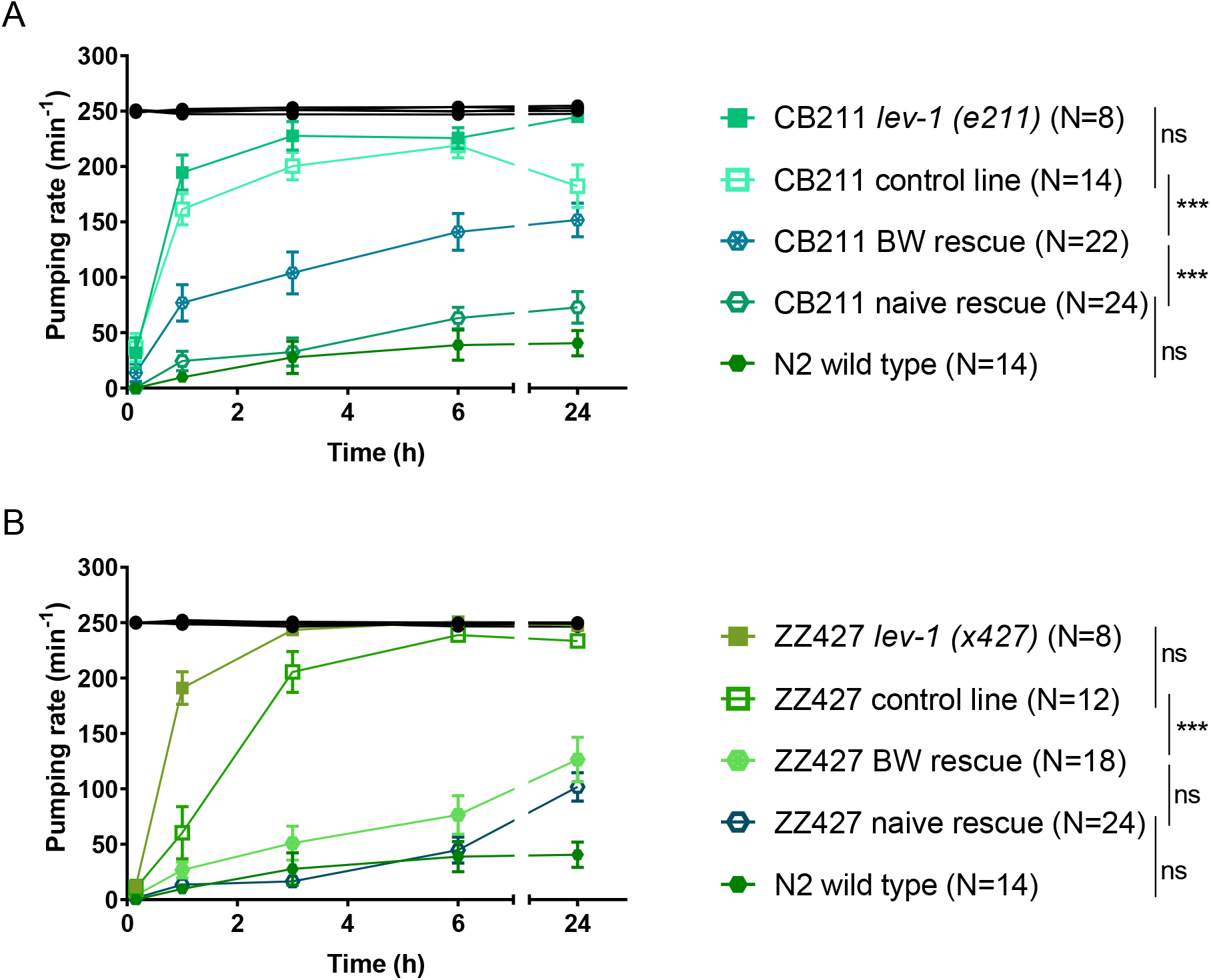
LEV-1 wild type expression in body wall muscles of *lev-1* mutant nematodes restores the levamisole induced inhibition of the pharyngeal function. A) Pharyngeal pumping in the absence (black) or presence (green) of 250 μM levamisole at different end-point times for N2, CB211 *lev-1 (e211)* mutant strain and transgenic lines expressing *lev-1* under either its own promoter (naïve rescue) or body wall muscle promoter (BW) into a CB211 background. Transgenic control lines were made by expressing GFP in coelomocytes of a *lev-1 (e211)*. Data are shown as mean ± SEM of the pumping rate of 14 worms in at least 7 different experiments for N2 wild type, 8 worms in at least 5 independent experiments for CB211 *lev-1 (e211)* mutant strain, 14 worms from two different lines in at least 4 independent experiments per line for control transgenic line, 22 worms from four different lines in at least 3 different experiments per line for body wall rescue transgenic lines and 24 worms from four different lines in at least 3 different experiments per line for naïve rescue lines. B) Pharyngeal pumping in the absence (black) or presence (green) of 250 μM levamisole at different end-point times for N2, ZZ427 *lev-1 (x427)* mutant strain and transgenic lines expressing *lev-1* under either its own promoter (naïve rescue) or body wall muscle promoter (BW) into a ZZ427 background. Transgenic control lines were made by expressing GFP in coelomocytes of a *lev-1 (x427)*. Data are shown as mean ± SEM of the pumping rate of 14 worms in at least 7 different experiments for N2 wild type, 8 worms in at least 5 independent experiments for ZZ427 *lev-1 (x427)* mutant strain, 12 worms from two different lines in at least 3 independent experiments per line for control transgenic line, 18 worms from three different lines in at least 3 different experiments per line for body wall rescue transgenic lines and 24 worms from four different lines in at least 3 different experiments per line for naïve rescue lines. ^ns^p>0.05; ***p<0.001 by two-way ANOVA test.

The naïve expression of *lev-1* rescued the wild type pharyngeal sensitivity to levamisole in the two *lev-1* deficient mutants tested (Fig. 5). Furthermore, the phenotype was partially rescued in the *lev-1 (e211)* mutant strain (Fig. 5A) and fully rescued in the *lev-1 (x427)* mutant strain (Fig. 5B) when the wild type version of the cDNA was selectively expressed in the body wall musculature under the control of the *myo-3* promoter.

These results indicate that the inhibition of the pharyngeal function by levamisole exposure is driven by the LEV-1 signalling at the body wall muscle.

### The pharyngeal sensitivity to levamisole is not mediated by a neuroendocrine signal

Pharyngeal pumping rate in mutants deficient in major transmitters was investigated after 6 hours of incubation with levamisole (Fig. 6), a time at which inhibition of pumping rate by this drug was dependent on the LEV-1 function at the body wall NMJ (Fig. 3 and 5). These strains included those deficient in *unc-25 (e156)*, *eat-4 (ky5)*, *cat-1 (ok411)*, *tph-1 (mg280)*, *tdc-1 (n3419)*, *tbh-1 (n3247)* and *egl-3 (n150)*.

**Figure 6:**
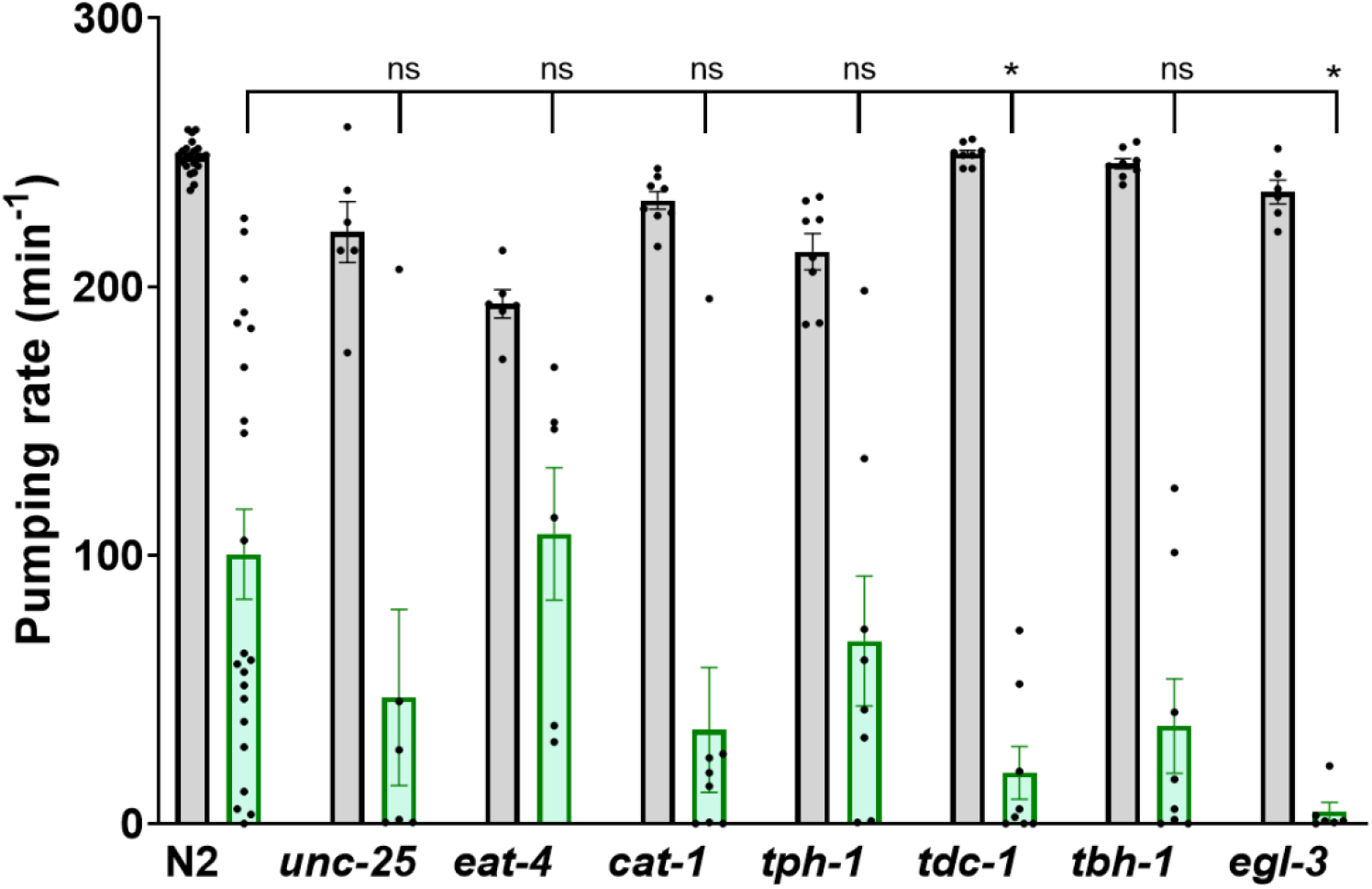
The inhibitory effect of levamisole in the pharyngeal function is independent of neurotransmitter or peptidergic signalling. Pumping rate of indicated neuromodulatory deficient mutants on food in the absence (grey column) or presence (green column) of 250 μM of levamisole after 6 hours incubation. Statistical analysis corresponds to the comparison between pumping rate on levamisole for N2 wild type and the different mutant strains. Data are shown as mean + SEM. ^ns^p>0.05; *p˂0.05 by two-way ANOVA test.

Nematodes deficient in the neurotransmitters GABA (*unc-25*), glutamate (*eat-4*) or the biogenic amines (*cat-1*) serotonin (*tph-1*), octopamine (*tbh-1*) or both, tyramine and octopamine, (*tdc-1*) exhibited a similar response to levamisole compared with the wild type nematodes. Since none of the mutant strains tested phenocopy the response observed in *lev-1* deficient worms, it indicates limited contribution of these major transmitter pathways to the levamisole-induced inhibition of the pharyngeal circuit or the underlying pharyngeal muscle pumping. (Fig. 6).

## Discussion

The screening performed in the present study was designed to identify molecular determinants that control the pharmacological inhibition of pumping during cholinergic hyperstimulation while comparing the cholinergic dependent intrinsic ability to respond to food (10). Following this, we clearly defined three groups of determinants.

### Physiological determinants of pharyngeal function

Although the pharyngeal muscle has two important cholinergic inputs in MC and M4 controlling its core function, there are additional cholinergic neurons of unknown function (17). Our study highlights an important class of mutants, including *eat-2* and *eat-18*, which are fundamental to sustain high pumping rate on food in physiological conditions (Fig. 1A). Indeed, our comparative approach strongly reinforces the critical role of the EAT-2/EAT-18 dependent receptor (21,25). The subunits of the nicotinic receptor ACR-6, ACR-10, DEG-3 and ACR-9 (29), the acetylcholine-gated chloride channel subunit ACC-3 (30) and the calcineurin CNB-1 (38) are included in this group (Fig. 7). This may have a value in better understanding additional roles of cholinergic signalling and associated receptors in feeding behaviour. However, further investigations will be required in order to identify the molecular pathways in which the physiological determinants of the feeding phenotype exert their function.

**Figure 7:**
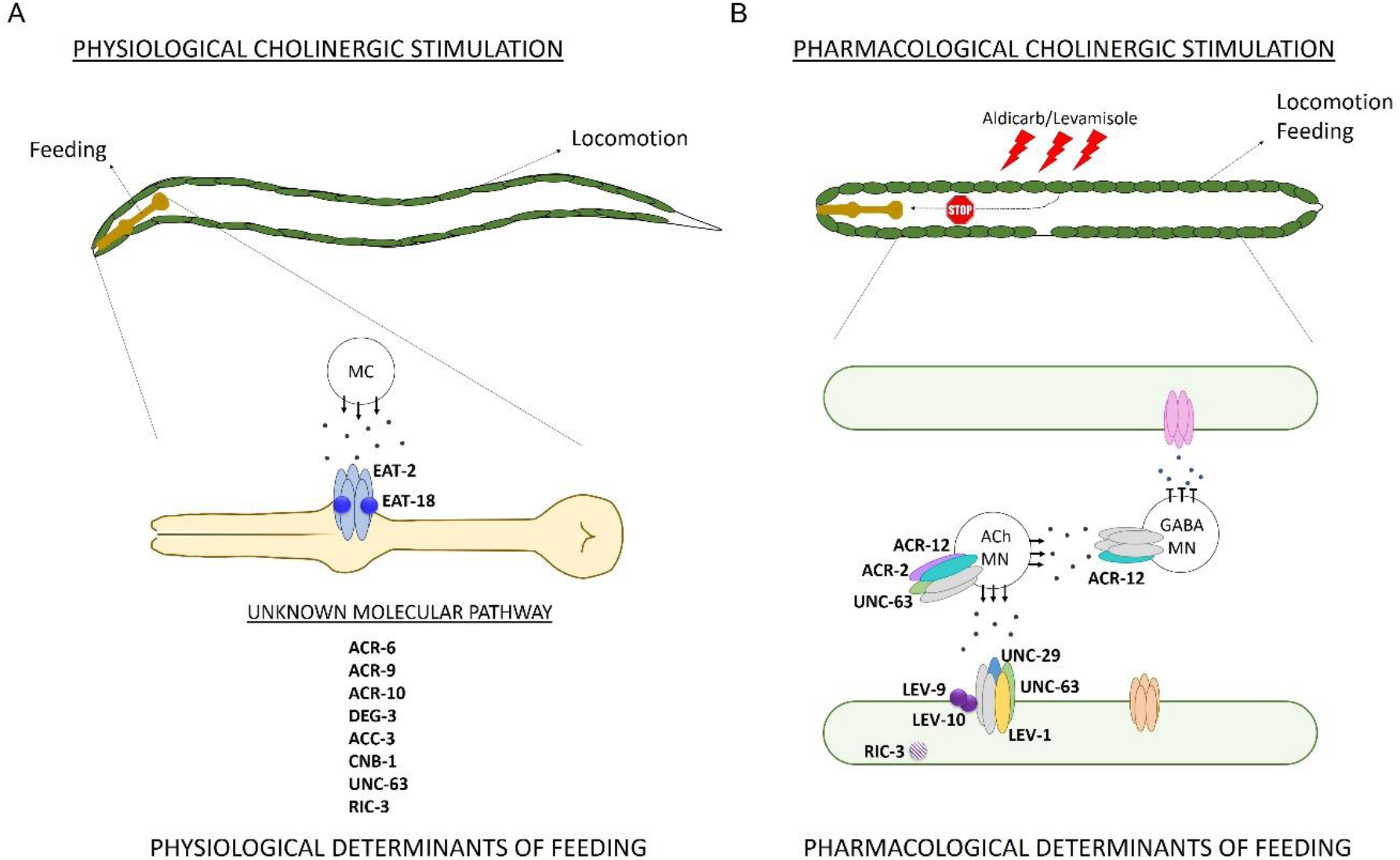
Physiological and pharmacological determinants of the feeding phenotype. The determinants of the pharyngeal function are distinct in the two contexts probed in this study. A) When nematodes are on food the cholinergic transmission stimulates pumping that underpins physiological feeding. The determinants of the pumping rate are EAT-2 and EAT-18, transducing the MC cholinergic signal in the pharyngeal muscle (21,25). The subunits of the nicotinic receptor ACR-6, ACR-9, ACR-10, DEG-3, ACC-3, UNC-63, the calcineurin subunit CNB-1 and the ancillary protein RIC-3 were identified as additional molecular determinants of pumping rate on food. B) The pharmacological overstimulation of the cholinergic pathway by aldicarb or levamisole drives a hypercontraction of the body wall muscle that imposes inhibition of the pumping rate. In this context, the determinants of the L-type receptor at the body wall musculature that underpin locomotion impart resistance to hypercontraction mediated inhibition of the feeding phenotype. These determinants are the subunits of the acetylcholine-gated ion channel UNC-63, UNC-29, LEV-1, ACR-2, ACR-12 and their auxiliary proteins LEV-9, LEV-10 and RIC-3.

Interestingly, none of the mutants included in this group displayed additional resistance to aldicarb-induced inhibition of the pharyngeal function (Fig. 1A). It indicates that the physiological determinants responsible for the essential control of the pharynx are quite distinct from those that drive the inhibition in the presence of aldicarb. Indeed, the distinct nature of mutants reinforces this proposition (Fig. 7).

### Pharmacological determinants of pharyngeal function

In the present study, we have used the aldicarb-induced paralysis protocol (26,27) to investigate determinants that regulate the pharyngeal pumping behaviour. However, this has highlighted a distinct non-pharyngeal modulation of drug-induced feeding inhibition. These investigations have been built on our previous observations indicating that aldicarb and other anti-cholinesterases cause a profound inhibition of the pharyngeal pumping (28). It was underpinned by a spastic paralysis of the radial muscles in the pharynx evidenced by an overt opening of the lumen (28). The prediction of our initial investigations was explained by assuming that aldicarb-dependent inhibition of acetylcholinesterase leads to an excess of input to the pharyngeal muscle from the two cholinergic motor neurons, MC and M4 (18,23,59,60). In contrast to this view, we demonstrated here that LEV-1 and other molecular components of the body wall NMJ are strong determinants of the aldicarb-induced inhibition of the pharyngeal function (Fig. 1B). This points to the pivotal role of the body wall muscle receptor in controlling locomotion and pharyngeal pumping in conditions where the pharmacological stimulation of the cholinergic signal causes an excitation of the musculature beyond the physiological levels (Fig. 7). This is reinforced by two observations: the failure to detect the *lev-1* expression in the pharynx (Fig. 4) and the tissue-specific rescue experiments in the *lev-1* mutant backgrounds (Fig. 5). The introduction of the wild type version of *lev-1* in the body wall muscle had a strong rescue effect of the levamisole-induced inhibition of pumping (Fig. 5). However, we note that *lev-1* is expressed more widely than body wall muscle. Therefore, the expression in the nerve cord and nerve ring could additionally contribute to the integrity of the response.

Overall, these results suggest that paralysis of locomotion by the signalling of the L-type receptor modulates pharyngeal function by inhibiting the pumping rate in that particular stress condition. Indeed, feeding can continue after the ablation of the pharyngeal neurons (9) but can be completely abolished by mechanical stimulation of the nematodes (15) or optical silencing of the body wall musculature (14). In the present study, we demonstrated that the pharmacological stimulation of the cholinergic signal at the body wall NMJ causes the reduction of the pharyngeal pumping (Fig. 7). This is a clear example of inter-tissue communication that is advantageous to the worm, allowing the coupling of two distinct functions. The mechanism underpinning the communication between the body wall and the pharyngeal NMJ is still unknown. Previously published observations highlighted the implication of dense core vesicle release as well as innexins as part of this mechanism (14). Using our pharmacological paradigm, we did not identify clear routes of chemical transmission responsible for the coupling between feeding and locomotion. Further investigations will be needed in order to underpin the signalling between the body wall and the pharyngeal circuits.

### Determinants playing a role in both scenarios

A final class of mutants that emerged from the screen includes *unc-63* and *ric-3* deficient strains. These two mutants are the only ones tested that exhibited a deficit in the pumping rate on food in the absence of aldicarb, and a resistance to the inhibition of pumping in the presence of the drug (Fig. 1C).

The central role of UNC-63 and RIC-3 in the composition and maturation of the body wall L-type receptor at the body wall muscle (39,50) explains why its deficiency impacts on the ability of aldicarb to induced the pharyngeal inhibition (Fig. 7). However, the fact that these mutants also impart the loss of the physiological pump rate on food suggests an under investigated role of UNC-63 and RIC-3 function within the pharyngeal circuit. This highlights paucity of understanding of the cholinergic determinants in pharyngeal function and how our screening approach may provide information about this in the future.

## Conclusion

In the *C. elegans* model organism, the ability of the pharynx to act as an interceptive cue for food to globally affect motility has been previously established (12). In the present work, we demonstrated that the pharmacological activation of the body wall circuit allows the distal inhibition of the pharyngeal pumping rate. This highlights a reverse route in which the tone of the musculature that controls locomotion impacts the circuit controlling the feeding behaviour. This fact provides insight into how the physiological state of one tissue can indirectly, but profoundly, impose control on distinct organs with an unrelated function. Acute regulation of pumping by the locomotory circuit has been noted (14,15), however, the advantages and mechanisms for allowing this remain to be resolved. In a wider sense, this kind of inter-tissue communication can report stress or disease in the whole organism’s physiology. In *C. elegans*, the hypercontraction of body wall muscles might act as an aversive cue that impacts in the feeding rate of the worm in a similar manner to the signals that are involved in disease in higher animals that impact on the appetite and feeding during cachexia (61).

## Experimental procedures

### *C. elegans* maintenance and strains

Nematodes were maintained at 20°C on NGM plates supplemented with *E. coli* OP50 strain as a source of food (62). *C. elegans* mutant strains are listed in Table 1 and were provided by Caenorhabditis Genetics Center (CGC) unless otherwise specified. Mutant strains EN39 *oig-4 (kr39)* II, EN300 *rsu-1 (kr300)* III and EN100 *molo-1 (kr100)* III were kindly provided by Jean-Louis Bessereau Lab (Institut NeuroMyoGène, France). ZZ427 *lev-1 (x427)* and transgenic lines AQ585 corresponding to N2; *Ex[Plev-1::gfp; rol-6]* genotype and AQ749 corresponding to ZZ427 *lev-1 (x427)* IV*; Is[Plev-1::lev-1::HA::gfp; rol-6]* genotype were kindly provided by William Schafer Lab (MRC Laboratory of Molecular Biology, UK). The transgenic lines GE24 *pha-1 (e2123)* III; *Ex[Punc-17::gfp; pha-1 (+)]* and N2; *Is[Peat-4::ChR2::mrfp]* were previously available in the laboratory stock.

The following transgenic lines were generated in this work: VLP1: CB211 *lev-1 (e211)* IV; *Ex[Punc-122::gfp]*; VLP2: CB211 *lev-1 (e211)* IV; *Ex[Punc-122::gfp; Pmyo-3::lev-1]*; VLP3: ZZ427 *lev-1 (x427)* IV; *Ex[Punc-122::gfp]*; VLP4: ZZ427 *lev-1 (x427)* IV; *Ex[Punc-122::gfp; Pmyo-3::lev-1]*; VLP5: CB211 *lev-1 (e211)* IV; *Ex[Punc-122::gfp; Plev-1::lev-1]*; VLP6: ZZ427 *lev-1 (x427)* IV; *Ex[Punc-122::gfp; Plev-1:lev-1]*.

### Generation of lev-1 rescue constructs

PCR amplifications were performed using Phusion High-Fidelity PCR Master Mix with HF Buffer (Thermo Fisher Scientific) following manufacturer instructions unless otherwise is specified.

PCR was used to amplify sequence for the *myo-3* promoter. 2.3 kb upstream of *myo-3* was amplified using the primers 5’ TCC<u>TCTAGA</u>TGGATCTAGTGGTCGTGG 3’ and 5’ ACC<u>AAGCTT</u>GGGCTGCAGGTCGGCT 3’ (58°C annealing temperature). This was subsequently cloned into pWormgate expression vector using the indicated restriction sites incorporated into the 5’ end of the oligonucleotides indicated above (HindIII/XbaI).

The primers 5’ AT<u>GCTAGC</u>TCTCATAACACTCAAGAAAACCCA 3’ and 5’ CCTCTATCCTCCACCACCTCCTAAC 3’ were used to amplify 3.536 kb of *lev-1 locus* corresponding to 3.5 kb upstream of the starting codon and 36 pb of exon one. PCR conditions for amplification were: initial 3 min at 98°C following of 34 cycles consisting of 1 min at 98°C, annealing 1 min at 57°C, extension 3:30 min at 72°C and a final extension 10 min at 72°C. Amplification product was cloned into pWormgate expression vector using the restriction site NheI underlined in forward primer and the naturally occurred XbaI restriction site 4 pb upstream of the *lev-1* starting codon.

cDNA of *lev-1* was amplified from a *C. elegans* cDNA library (OriGene) using 5’ AGAGAGAATGATGTTAGGAGG 3’ and 5’ AGTTGAAAATGAAAGAATAATGG 3’ (55°C annealing temperature) forward and reverse primers, respectively. The PCR product was subcloned into pCR8/GW/TOPO following manufacturer protocol and subsequently cloned into pWormgate plasmid containing either *Pmyo-3* or *Plev-1* in order to generate *Pmyo-3::lev-1* and *Plev-1::lev-1* plasmids, respectively. The sequence of the plasmids was validated by Sanger sequencing before microinjection.

### Generation of transgenic lines

The marker plasmid *Punc-122::gfp* was kindly provided by Antonio Miranda Lab (Instituto de Biomedicina de Sevilla, Spain). It drives the expression of GFP specifically in coelomocytes of *C. elegans* (Miyabayashi, Palfreyman et al. 1999).

The microinjection procedure was performed as previously described (63). A concentration of 50 ng/μl of the marker plasmid *Punc-122::gfp* was injected into one day old adults of the CB211 *lev-1 (e211)* IV and ZZ427 *lev-1 (x427)* IV mutant background to generate the transgenic strains VLP1 and VLP3, respectively. A mixture of 50 ng/μl of *Punc-122::gfp* plasmid and 50 ng/μl of *Pmyo-3::lev-1* plasmid was microinjected into adults of CB211 and ZZ427 strains to produce the transgenic lines VLP2 and VLP4, respectively. Finally, transgenic strains VLP5 and VLP6 were generated by microinjecting adults of the *lev-1 (e211)* and *lev-1 (x427)* mutant backgrounds with a mixture of 50 ng/μl of *Punc-122::gfp* plasmid and 50 ng/μl of *Plev-1::lev-1* plasmid, respectively.

The genotype of CB211 and ZZ427 strains was authenticated by PCR amplification of the *lev-1* gene and subsequent sequencing of the PCR product before microinjection was carried out.

### Plate husbandry

Aldicarb and levamisole hydrochloride (Merck) were dissolved in 70% ethanol and water, respectively. The stock drugs were kept at 4°C and used within a month or discarded.

Behavioural experiments were performed at room temperature (20°C) in 6-well plates that were prepared the day before of each experiment. Drug-containing plates were made by adding a 1:1000 aliquot of the concentrated stock to molten tempered NGM agar to give the indicated concentration of aldicarb (500 μM) and levamisole (10 μM to 500 μM). For aldicarb control plates, a similar aliquot of 70% ethanol was added to the molten agar. The final concentration of ethanol for control and aldicarb-containing plates was 0.07%. This concentration of ethanol did not affect any of the behavioural tests performed in this work (data not shown).

For protracted intoxication experiments, 50 μl of OP50 bacteria culture OD_600_ 1 was pipetted onto the solidified NGM assay plates containing either drug or vehicle. For the first 10 min of exposure to levamisole (early intoxication experiment), assay plates were seeded with 100 μl of OP50 bacteria culture of one OD_600_ that was spread evenly over the complete surface of the NGM agar. After seeding, plates were left in the laminar flow hood for one hour to facilitate drying of the bacterial lawn. 6-well plates containing either levamisole or aldicarb with bacterial lawn were then stored in dark at 4°C until next day. Assay plates were incubated at room temperature for at least 30 min before starting the experiment. Contrary to assay plates with other cholinergic drugs (64), we did not observe any difference in the density or integrity of the bacterial lawn between control and drugged plates.

### Behavioural observations

A pharyngeal pump as a cycle consists of contraction-relaxation of the terminal bulb in the pharyngeal muscle. Each pump was discerned by the backward-forward movement of the grinder structure in the terminal bulb. The quantification was made visually under a binocular dissecting microscope Nikon SMZ800 (x60 magnification) using a clicker counter.

For protracted intoxication experiments, synchronized nematodes one day older than L4 stage (L4 + 1) were transferred onto the assay plates and the pumping measured after 24 hours for aldicarb intoxication assays and after 10 min, 1, 3, 6 and 24 hours for levamisole intoxication experiments. Nematodes that left the patch of food during the experiment were picked back to the bacterial lawn and the pumping rate was scored after waiting for between 10-15 min.

For the first 10 min of exposure to levamisole, synchronized (L4 + 1) adults were picked onto either control or levamisole-containing plates. The delay between each pump was scored for the consecutive 10 min straight after transferring each worm using Countdown Timer tool from www.WormWeb.org website. It was then translated into pumping rate per second.

### Pharynx dissection procedure

Dissection of the pharynxes was performed according to previously published methods (65). Young adult (L4 + 1) worms were placed into dissection plates containing 3 ml of Dent’s solution (glucose 10 mM, HEPES 10 mM, NaCl 140 mM, KCl 6 mM, CaCl_2_ 3 mM, MgCl_2_ 1 mM) supplemented with 0.2% bovine serum albumin (Merck). Dishes were incubated at 4°C for 5 min to reduce the thrashing activity of the nematodes and then placed under a binocular microscope Nikon SMZ800. The lips of the worms were dissected from the rest of the body by making an incision with a surgical scalpel blade. Due to the internal pressure of the inside organs of the worm, the content is ejected outside the cuticle of the nematode leaving the pharynx and its embedded circuit exposed. When the terminal bulb was clearly observed outside the cuticle, a second incision was made at the pharyngeal-intestinal valve to isolate the pharynx from the rest of the intestines (Fig. 4). Pharynxes lacking more than half of the procorpus after dissecting were not considered for either imaging or for RT-PCR.

### Differential interference contrast (DIC) and fluorescence imaging of pharyngeal structure and transgene expression

Isolated pharynxes were removed from dissection dishes using non-sticky tips within 10 μl of solution. Fat and debris were carefully removed from the pharynxes by two sequential transfers, in a volume of 10 μl, through two changes of 3 ml Dent’s 0.2% BSA media. After washing, the pharynxes were placed on a thin pad of 2% agarose previously deposited and solidified on a microscope slide. A 24×24 mm coverslip was gently located on top before observations were made. Objectives of 10x/0.30, 60x A/1.40 (oil) and 100x A/1.40 (oil) fitted in a Nikon Eclipse (E800) microscope were used to collect images through both DIC and epifluorescence filters. A Nikon C-SHG1 high pressure mercury lamp was used for illumination in fluorescence micrographs.

Images were acquired through a Hamamatsu Photonics camera software, and were cropped to size, assembled, and processed using Abode PhotoShop^®^ (Adobe Systems) and ImageJ (NIH) software.

Two transgenic strains harbouring *Punc-17::gfp* and *Peat-4::rfp* were used as control of the dissection procedure. They are both transcriptional reporters of cholinergic and glutamatergic neurons, respectively. Isolated pharynxes from the two transgenic strains were isolated and imaged following the previously explained protocol. Fluorescent from distinct neurons was observed upon the dissection procedure (data not shown) indicating the pharyngeal neurons were preserved in the isolated pharynx preparations.

### RT-PCR from whole worms and isolated pharynxes

RT-PCR for single worm and isolated pharynxes was performed following previously published method with modifications (66). A single worm or isolated pharynxes were placed into 1 μl of worm lysis buffer containing a final concentration of 5 mM Tris pH8, 0.25 mM EDTA, 0.5% Triton X100, 0.5% Tween 20 and 1 mg/ml proteinase K. After a brief centrifugation, the mixture containing either the single worm or the isolated pharynxes, was incubated at 65°C for 10 min and at 85°C for 1 min into a T100™ thermocycler (Bio-Rad). The heated lysate was subsequently used in cDNA synthesis with SuperScript™ III Reverse Transcriptase kit in a total volume of 20 μl following manufacturer protocol (Invitrogen™). 5 μl of the resulting cDNA was used for a final volume of 20 μl PCR with indicated oligo primers. PCR amplifications were performed employing Phusion High-Fidelity PCR Master Mix with HF Buffer (Thermo Scientific™) following manufacturer’s recommendations.

The primers used for the PCR extensions were 5’ GGACAGGGAGCCGAGAAGAC 3’ and 5’ GAAGCATCGTTAAGGAAAGTCAGG 3’ (64°C annealing temperature) for *myo-2*; 5’ GTGAATAGTCAGTTGGTGATGG 3’ and 5’ TGCGAAAATAAGTGCTGTGGTG 3’ (66°C annealing temperature) for *eat-2*; 5’ CTGCTCGTCCTTTTGATTCC 3’ and 5’- GTTTCCCTTCACAGTTACCAC -3’ (58°C annealing temperature) for *myo-3*; 5’ ATGTTAGGAGGTGGTGGAGG 3’ and 5’ GTTGAACGAGAGAGTTGTATCC 3’ (66°C annealing temperature) for *lev-1*.

### Statistical analysis

The collection of the data was performed blind, so the experimenter was unaware of the genotype and the drug present, absent or concentration tested in each trial.

Data were analysed using GraphPad Prism 8 and are displayed as mean ± SEM. Statistical significance was assessed using two-way ANOVA followed by post hoc analysis with Bonferroni corrections where applicable. This post hoc test was selected among others to avoid false positives. The sample size N of each experiment is specified in the corresponding figure.

## Acknowledgements

We thank Dr Jean-Louis Bessereau, Dr Denise Walker and Dr William Schafer for sharing strains; Dr Antonio Miranda-Vizuete for sharing *Punc-122::gfp* marker plasmid.

We thank Emeritus Professor Robert Walker, Dr Fernando Calahorro, Helena Rawsthorne-Manning and Johanna Haszczyn for critical reading of the manuscript and for detailed comments.

Additional *C. elegans* strains were provided by the CGC, which is funded by NIH Office of Research Infrastructure Programs (P40 OD010440).

## Author contributions

**Patricia G. Izquierdo**: Conceptualization, Data curation, Formal analysis, Investigation, Methodology, Validation, Visualization, Roles/Writing - original draft. **Thibana Thisainathan**: Investigation. **James Atkins**: Investigation. **Christian J. Lewis**: Investigation. **John Tattersall**: Conceptualization, Funding acquisition, Methodology, Supervision, Writing - review & editing. **Christopher Green**: Conceptualization, Funding acquisition, Methodology, Supervision, Writing - review & editing. **Lindy Holden-Dye**: Conceptualization, Funding acquisition, Methodology, Supervision, Writing - review & editing. **Vincent O’Connor**: Conceptualization, Funding acquisition, Methodology, Supervision, Writing - review & editing.

